# Body mass decline in a Mediterranean community of solitary bees supports the size shrinking effect of climatic warming

**DOI:** 10.1101/2022.11.03.515021

**Authors:** Carlos M. Herrera, Alejandro Núñez, Javier Valverde, Conchita Alonso

## Abstract

The long-known, widely documented inverse relationship between body size and environmental temperature (“temperature-size rule”) has recently led to predictions of body size decline following current climatic warming (“size shrinking effect”). For keystone pollinators such as wild bees, body shrinking in response to warming can have pervasive effects on pollination processes, but there is still little evidence of the phenomenon because adequate tests require controlling for climate-linked confounding factors (e.g., urbanization, land use change). This paper tests the shrinking effect in a diverse community of solitary bees from well-preserved habitats in the core of a large nature reserve undergoing fast climatic warming but not disturbances or habitat changes. Long-term variation in mean body mass was evaluated using data from 1,186 individual bees (108 species, 25 genera, 6 families) sampled over 1990-2022. Climate of the region warmed at a fast rate during this period and changes in bee body mass verified expectations from the size shrinking effect. Mean individual body mass of the regional community of solitary bees declined significantly, shrinking being particularly intense for female individuals and large-bodied species. As a consequence, the pollination and mating systems of many bee-pollinated plants in the region are likely undergoing important alterations.

## Introduction

Intra and interspecific variation in individual body size is often predictably related to environmental temperature. Early recognition of such relationships led to the formulation of some classical ecological rules such as Bergmann’s, Allen’s or the “temperature-size rule” (Ray 1960, Atkinson 1994). The temperature-size rule, a “widely documented and poorly understood pattern” (Klok and Harrison 2013), predicts an inverse relationship across conspecific individuals between body size and the environmental temperature experienced during development (Atkinson 1994, Forster and Hirst 2012, Klok and Harrison 2013). While the proximate and ultimate causes of the temperature-size rule are still far from being fully understood (Walters and Hassall 2006, Verberk et al. 2021), it is a matter of fact that the rule has proven true in the majority of ectothermic organisms where it has been investigated (Atkinson 1994, Klok and Harrison 2013). Such predictable relationship between body size and environmental temperature has recently acquired particular relevance in current ecological scenarios affected by climate warming, motivating predictions of reduced body size in response to rising temperatures (“size shrinking effect”; Sheridan and Bickford 2011, Ohlberger 2013, Verberk et al. 2021).

Given the many life history, ecological and evolutionary consequences of individual body size (Bonner 2006), the latter’s predicted reduction attached to climatic warming can trigger a complex cascade of effects whose details will depend on the ecological functionality of the organisms under consideration. In the case of keystone pollinators such as wild bees, body shrinking can alter crucial aspects of the pollination process such as foraging range, pollen load size and diversity, and pollen carryover, all of which are related to body size (Greenleaf et al. 2007, Cullen et al. 2021, Földesi et al. 2021). Limited evidence suggests that recent changes in bee sizes conform to expectations from the temperature-size rule, but studies so far refer to few species, consider linear measurements as a proxy for size rather than actual body mass, and the putative effects of temperature can be confounded with those of other factors correlated with climate change such as urbanization, land use changes or habitat fragmentation (Oliveira et al. 2016, Grab et al. 2019, Kelemen and Rehan 2021, Suni and Dela Cruz 2021, Garlin et al. 2022). In this paper we test size shrinking effects expected from the temperature-size rule by examining long-term changes in individual body mass in a diverse community of solitary bees sampled over three decades in Mediterranean habitats from a large protected area. Our study region is undergoing climatic warming but not disturbances or habitat changes that could confound the effects of the warming climate.

## Materials and methods

As part of other studies (Herrera 1995, 1997, Herrera et al. 2022), a total of 1,186 solitary bees from 108 species, 25 genera and six families (Andrenidae, Apidae, Colletidae, Halictidae, Megachilidae, Melittidae; see Appendix S1: Table S1, for species and sample sizes) were hand-netted in the field during 1990-1997 (“old period” hereafter; *N* = 472 individuals, 46 species) and 2022 (“recent period”; *N* = 714 individuals, 99 species), in 47 localities located in the core area of the Sierras de Cazorla, Segura y Las Villas Natural Park, a 2,140 km^2^ protected area in Jaén Province, southeastern Spain. The species sampled represented ∼30% of all species of bees occurring in the region, and mostly belonged to the genera *Andrena* (643 individuals, 32 species), *Colletes* (82, 4), *Anthophora* (81, 10), *Osmia* (72, 4), *Anthidium* (64, 2) and *Xylocopa* (51, 3) (Appendix S1: Table S1). A map of sampling locations is shown in Appendix S2: Figure S1. The sampled area is characterized by their well-preserved habitats and outstanding biological diversity. Our sampling sites were located at elevations between 740–2107 m, and did not undergo any obvious disturbance or habitat changes over the 30-yr period considered here. Netted bees were placed individually into sealed microcentrifuge tubes, kept in the dark in a portable refrigerator, and brought to the laboratory within a few hours, where they were weighed to the nearest 0.1 mg using always the same Sartorius 1602MP8 analytical balance.

Recent increase in environmental temperature has been well documented for the southeastern Iberian Peninsula (Fernández-Montes and Rodrigo 2015, Gonzalez-Hidalgo et al. 2015). To corroborate this trend at the spatial and temporal scale of this study, daily maximum and minimum temperature data for 2000-2022 were gathered for 10 weather stations 15-60 km away from our bee sampling sites (Red de Información Agroclimática de Andalucía; https://www.juntadeandalucia.es/agriculturaypesca/ifapa/riaweb/web/; last accessed 23 October 2022). Relevant details, including a location map in relation to bee sampling sites are shown in Appendix S2: Figure S1, Table S1.

A similar analytical framework based on fitting linear mixed effects models to the data (“mixed models” hereafter; Bolker 2015, Harrison et al. 2018) was adopted to test for supra-annual trends in both environmental temperature and body mass of solitary bees. In the case of temperature, random intercepts models were fit to daily maximum and daily minimum temperature data for every weather station. These models had year as the single fixed effect (treated as a continuous, numerical variable) and Julian date (= days since 1st January, treated as an unordered factor with 365 levels) as random effect. Fixed-effect parameters obtained from these models provided estimates of the rate of change of annual means for daily maximum and minimum temperatures. For bee body mass, a random slopes–random intercepts model was fit to the individual mass data (log_10_ transformed), with year of capture (centered and scaled to facilitate model convergence and interpretation of fixed effects and interactions; Harrison et al. 2018) as fixed effect and bee species as random effect. Since sexual size dimorphism is frequent in solitary bees (Danforth et al. 2019), sex and its interaction with year of capture were included in the model as fixed-effect covariates. As a significant sex x year interaction was found, model estimates for the annual rate of change of body mass were computed separately for males and females. Preliminary analyses including sampling date (days from 1st January) and altitude of sampling site as additional fixed-effect covariates did not improve model fit significantly. These two variables were omitted from the final analyses for simplicity.

The random slopes from our mixed model reflected variation among levels of the random effect (bee species) in the effect of the predictor (year) on the response variable (body mass). We evaluated whether body mass decline rates were related to body mass (Klok and Harrison 2913) by testing, across species, the relationship between estimated slopes and mean body mass. To verify the theoretically expected robustness of fixed-effect parameter estimates to the small sample sizes of many species (i.e., sparsely sampled random effect levels; Bolker 2015), models were fit to the whole 108 species dataset and only to the subset of 33 species which were sampled in both the old and recent periods. Numerical results of the two analyses were virtually identical, so only those for the whole sample will be reported.

All statistical analyses were carried out using the R environment (R Core Team 2022). Mixed models were fit with the lmer function in the lme4 package (Bates et al. 2015). Model adequacy was assessed using the check_model function in the performance package (Lüdecke et al. 2021). Model fit to data was assessed with the conditional and marginal *R*^2^ for each model (Nakagawa and Schielzeth 2013), obtained with the r2_nakagawa function from package performance. Confidence intervals of fixed-effect parameters obtained by bootstrapping (function bootstrap_model, package parameters; Lüdecke et al. 2020) and Wald Chi-square tests (Anova function, car package; Fox and Weisberg 2019), were used to assess the statistical significance of fixed effects. *P* values for multiple tests of the same hypothesis were corrected using the Benjamini-Hochberg procedure (p.adjust function, stats package). The function ggpredict from the ggeffects package (Lüdecke 2018) was used to compute marginal effects of year on mean temperature and mean bee body mass.

## Results

The study area warmed significantly over the period 2000-2022. This trend was mainly due to a steep increase in yearly means of daily maximum temperatures, which took place consistently at all weather stations (Fig. 1). All mixed models fit to daily maximum temperature data yielded statistically significant, positive temperature/year relationships, with model-estimated slopes averaging +0.069 ºC/year and ranging between +0.042 and +0.12 ºC/year depending on the station (see Appendix S2: Table S1, for parameter estimates and confidence intervals). Increases in mean daily minimum temperature also took place, but were less marked (Fig. 1). Statistically significant relationships between daily minimum temperature and year occurred in only six weather stations, and in these cases the estimated slopes averaged +0.036 ºC/year and ranged between +0.012 and +0.088 ºC/year (Appendix S2: Table S1).

**Fig. 1.**
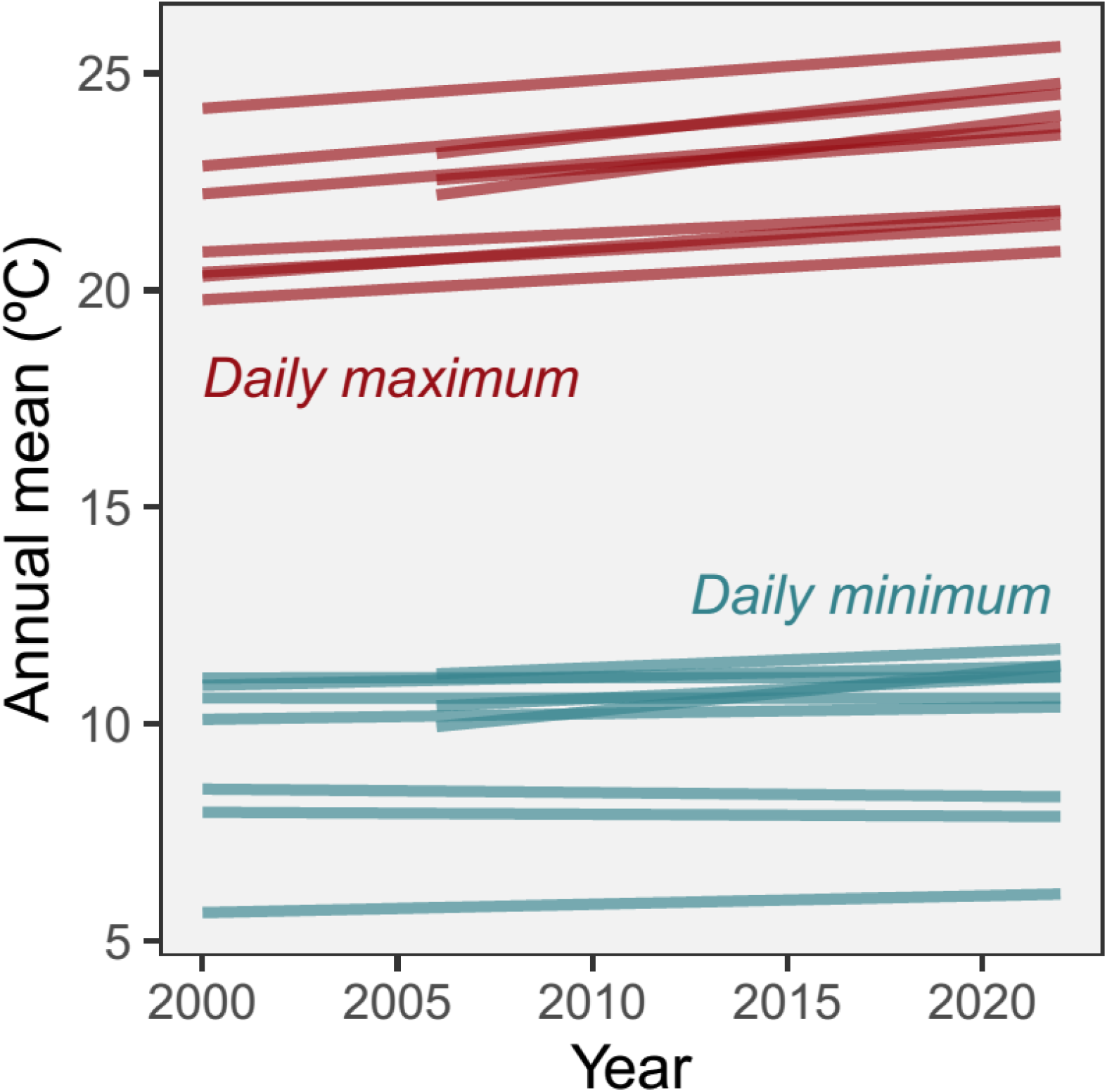
Linear trends in annual means for daily maximum and daily minimum temperatures for 10 weather stations near and around the bee sampling sites (see Appendix S2: Figure S1 for a map). Each line is the prediction obtained from the linear mixed model fit to daily temperature data for one station, with year as the single fixed effect and Julian date (unordered, qualitative factor) as random effect. A summary of analytical results is given in Appendix S2: Table S1.

Mean individual body mass of the community of solitary bees declined over the 1990-2022 sampling period. The mixed model fit reasonably well to the (log_10_ transformed) individual body mass data (conditional and marginal *R*^2^ = 0.954 and 0.109, respectively) and underlying model assumptions were reasonably supported (Appendix S3: Figure S1). There was a negative effect of year on body mass after statistically accounting for sexual size dimorphism and interspecific variation in intercepts and slopes of the mass/year relationship (Table 1). Sexes differed in their declining trends, as denoted by the significant year x sex interaction effect on body mass (Table 1). The decline was considerably steeper for females (slope ± SE = -0.0398 ± 0.0064) than for males (−0.0126 ± 0.0068). Because of the transformations applied to the data, these slopes represent changes in log_10_(mass) per year standard deviation unit and thus reflect multiplicative rather than additive changes. Back-transforming estimated slopes to original scales of measurement yields average body mass declines of 0.71%·year^-1^ and 0.23%·year^-1^, or an average cumulative reduction of ∼15 mg and ∼3 mg per individual bee from 1990 to 2022 for females and males, respectively (Fig. 2). Bee body mass decline at the community level revealed by the linear mixed model was also obvious in paired within-species comparisons for the subset of 33 species which were sampled in both the old and recent periods (Appendix S4: Figure 1) (*P* = 0.0004 and <0.0001 for males and females, respectively, permutation test).

**Table 1.** Summary of results of the linear mixed model fit to body mass data for individual bees sampled between 1990–2022 (*N* = 1,182 individuals, 105 species). Estimated model parameters and variances are expressed in the transformed scales (body mass log_10_ transformed, years scaled to mean = 0 and standard deviation = 1; see text for interpretations in the original scales). Checks of main model assumptions (linear relationship, homogeneity of variance, noncollinearity, normality of residuals, normality of random effects) are shown in Appendix S3: Figure S1.

| Model component | Parameter estimate<br>[95% confidence interval] | Chi-<br>squared | *P*-value | Variance<br>[95% confidence interval] |
| --- | --- | --- | --- | --- |
| Fixed effects |  |  |  |  |
| Year | -0.0398 [-0.0511, -0.0277] | 27.9 | 1.3e-07 |  |
| Sex (Male) | -0.2897 [-0.3041, -0.2737] | 1390.4 | <2.2e-16 |  |
| Year x Sex (Male) | 0.0272 [0.0127, 0.0417] | 14.3 | 0.00015 |  |
| Random effects |  |  |  |  |
| Bee species (intercept) |  |  |  | 0.1687 [0.1253, 0.2218] |
| Bee species (slopes) |  |  |  | 0.00047 [0.00011, 0.00099] |

**Fig. 2.**
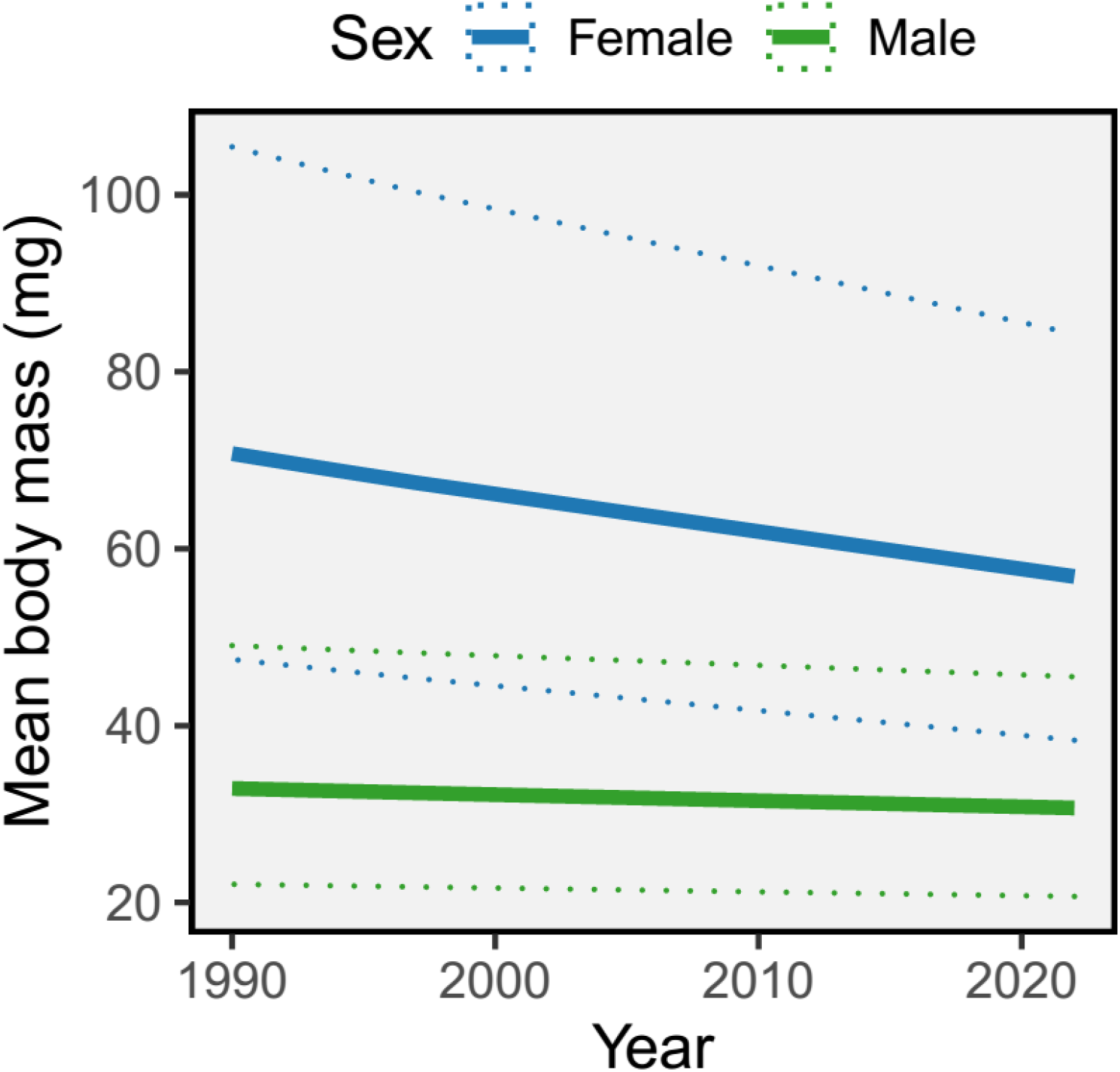
Mean estimated marginal effects of year on individual body mass of female and male bees (solid lines), as predicted from a mixed model with body mass as response variable, year, sex and their interaction as fixed effects, and bee species as random effect. Dotted lines are the 90% confidence intervals around estimated means, see Table 1 for numerical results for the model. The original analysis was conducted on transformed variables, and the mean marginal effects shown here are the back-transforms to the original measurement scales.

Yearly rate of body mass decline differed among bee species, as denoted by the non-zero variance component associated with random slopes (Table 1). Estimated slopes were negative for all species, encompassed two orders of magnitude (from -0.0801 to -0.00050), and were inversely correlated with log_10_(mean body mass) (*r* = -0.700, *N* = 108, *P* = 2.2e-16). This result was not an statistical artifact because intercepts and slopes were uncorrelated (*r* = -0.39, 95% confidence interval = -0.96, +0.16). The heavier a bee species, the steeper the proportional declining rate of body mass with years, and this pattern occurred consistently across all families represented in our sample (Fig. 3). While for the smallest bees (< 30 mg) the estimated mass declining rate ranged between -0.35 and -0.70%·year^-1^, the corresponding figures for the largest bees (>100 mg) were -0.97 and -1.40%·year^-1^.

**Fig. 3.**
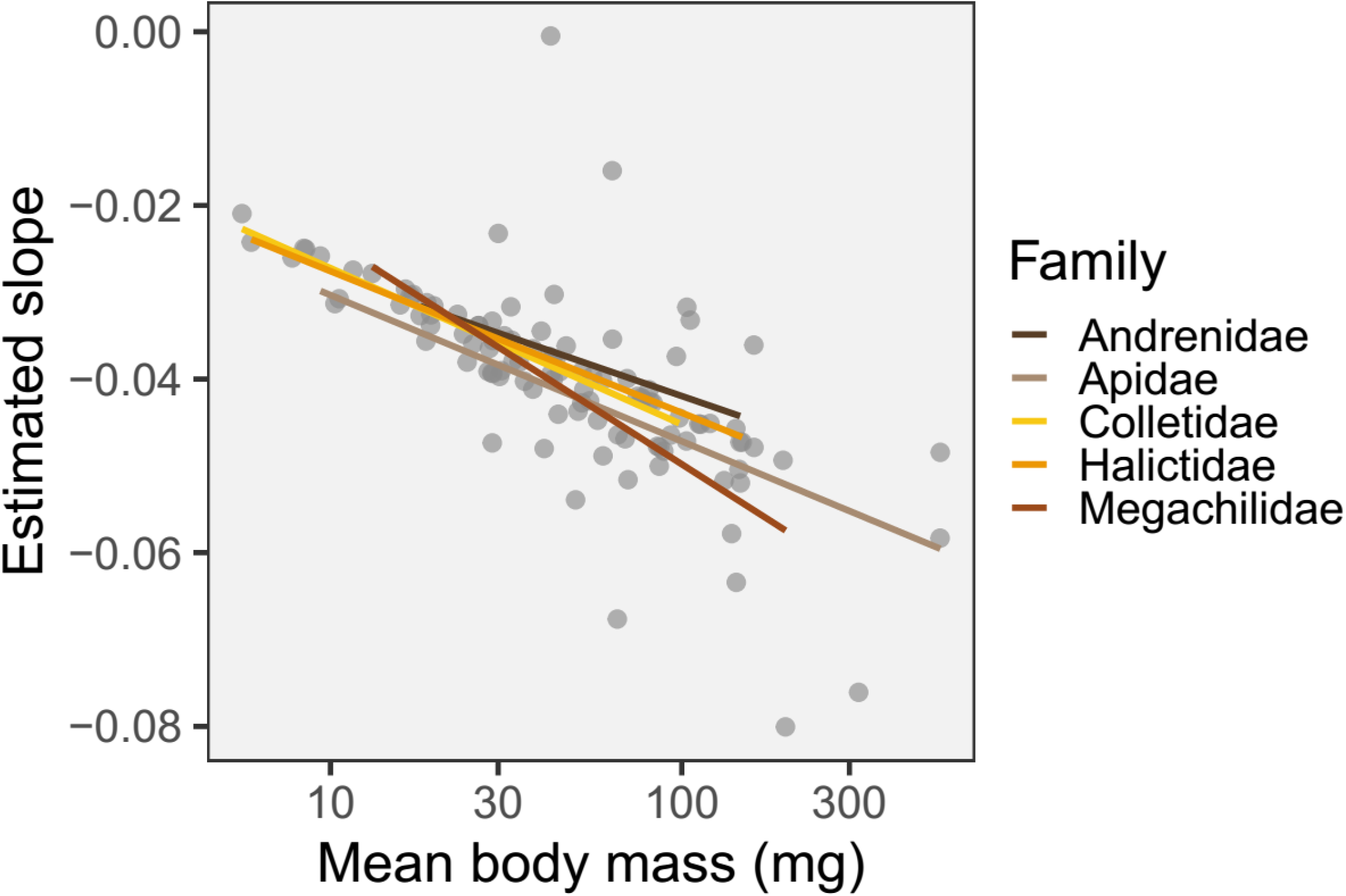
Inverse relationship between the slopes of log_10_(body mass)/year relationships (estimated from the mixed model fit to the data, Table 1) and the mean body mass of individual bee species (*N* = 105). Lines are regressions obtained separately for the different bee families.

## Discussion

The climate of our study area warmed at a fast rate during the last few decades, as shown by increasing mean daily maximum and, to lesser extent, minimum temperatures, thus confirming at the reduced spatial scale of our study the general trend for southeastern Spain (see references in Introduction). In agreement with expectations from the size shrinking effect, the warming climatic trend was concomitant with a decline in the mean individual body mass of the regional community of solitary bees, an effect which was strongest among females and largest-bodied species. Since bees were sampled in a well-preserved, protected area located ≥ 10 km away from urbanized or agricultural land, body size reduction can be parsimoniously attributed to the effects of climate warming. These results provide the first direct evidence to date of size shrinking in a diverse wild bee community based on body mass data rather than size metrics based on linear measurements of body parts, which are poor predictors of intraspecific variation in body mass (Kendall et al. 2019) and can produce biased results because of heterogeneous responses of body parts to temperature (Ray 1960, Klok and Harrison 2013). Adding robustness to our results is the treatment of the different bee species as levels of a random effect. Mixed models allow to make inferences that apply to different populations of effects, or “inference spaces” (Schabenberger and Pierce 2002). In the context of this study, the whole regional community of solitary bees represents the “broad inference space” and our conclusions refer specifically to that space, not just the 108 species sampled. This means that model parameter estimates for fixed effects refer to temporal changes in mean individual body mass for the solitary bee community as a whole (see Herrera 2019, for further discussion on the value of treating species as random effect levels when investigating community-level trends).

As it often happens with natural patterns conforming to the temperature-size rule (Klok and Harrison 2013, Verberk et al. 2021), the possible mechanism(s) responsible for the decline in body mass of solitary bees documented in this paper can only be tentatively suggested. The fact that all species consistently exhibited negative body mass slopes suggests that the ultimate cause of size reduction was universal enough as to affect all species similarly, irrespective of habitat type or nesting habits (e.g., ground-*vs*. hole-nesting). Experimental studies have documented inverse effects on body size of the temperature experienced during larval growth for some genera or species included in this study (e.g., *Osmia bicornis, Lasioglossum*; Kamm 1974, Radmacher and Strohm 2010, Kierat et al. 2017). These findings are in line with the size-temperature rule, and point to ubiquitous direct effects of rising temperatures on body shrinking of solitary bees in our study region. Nevertheless, additional indirect effects on adult body size mediated by temperature-dependent reductions in the quantity and quality of food provisions available to the larvae (Chole et al. 2019) can not be ruled out.

Larger bee species and, within species, the larger-bodied females, experienced the largest proportional reductions in body mass. Regardless of the factors responsible for this differential response, which cannot be addressed with the data available, sex- and size-dependent reduction in bee body size has three remarkable implications. Firstly, sex-dependent reduction rate of body size has produced a noticeable reduction in the extent of sexual size dimorphism (female size advantage declined from 38 mg to 26 mg on average; Fig. 2). Size differences between males and females of the same species drive key biological processes, and decreased sexual dimorphism can distort patterns of sexual selection and mating behavior and drive populations away from optimal sex investment ratios shaped by natural selection (Helms 1994, Alcock 2013), with unpredictable consequences for long term survival of populations. Secondly, the close relationship linking female body size and fecundity in insects (Honek 1993) suggests that per-capita fecundity of the largest species of solitary bees has probably declined substantially in our study region over the last few decades, which could account for the noticeable rarefaction of some large-sized species (e.g., *Andrena assimilis, A. labialis, A. thoracica, Xylocopa cantabrita*; C. M. Herrera, *unpublished data*). And thirdly, larger species and, within species, the larger female individuals tend to forage over wider areas, perform more and more effective pollinations, and deposit larger and more diverse pollen loads with a larger carryover (Herrera 1987, Greenleaf et al. 2007, Cullen et al. 2021, Földesi et al. 2021). The fact that female and larger bees are experiencing the steepest declines in body mass suggests that the pollination and mating systems of many bee-pollinated plants of our study region can be silently undergoing far-reaching changes.

## Acknowledgments

We are grateful to Consejería de Medio Ambiente (Junta de Andalucía) for permission to work in Sierras de Cazorla, Segura y Las Villas Natural Park, and Mónica Medrano and Juli Pausas for suggestions. Work partly funded by Ministerio de Ciencia e Innovación through European Regional Development Fund (SUMHAL, LIFEWATCH-2019-09-CSIC-13) and Consejería de Transformación Económica, Industria, Conocimiento y Universidades, Junta de Andalucía (P18-FR-4413).

## Author contributions

C. M. H. conceptualized the study, analyzed the data and led writing. A. N. identified bee specimens. C. A. provided financial support and project management. All authors participated in field and laboratory work, contributed to interpretation of results, edited and revised manuscript drafts, and approved the final version.

