## Appendix S1 for "Body mass decline in a Mediterranean community of solitary bees supports the size shrinking effect of climatic warming"

**Appendix S1. Bee species list and sample sizes**

TABLE S1. List of the 108 species of solitary bees sampled and number of individuals weighed per species and year.

| | Number of individuals weighed | | | | | | | |
|---|---|---|---|---|---|---|---|---|
| Species | 1990 | 1991 | 1992 | 1993 | 1994 | 1995 | 1997 | 2022 |
| *Amegilla albigena* | 0 | 0 | 0 | 0 | 0 | 0 | 0 | 13 |
| *Amegilla quadrifasciata* | 0 | 0 | 1 | 0 | 0 | 0 | 0 | 6 |
| *Andrena asperrima* | 0 | 0 | 0 | 0 | 0 | 0 | 0 | 2 |
| *Andrena assimilis* | 0 | 0 | 0 | 0 | 7 | 23 | 0 | 19 |
| *Andrena baetica* | 0 | 0 | 0 | 0 | 0 | 0 | 0 | 2 |
| *Andrena bicolor* | 0 | 0 | 14 | 3 | 5 | 3 | 0 | 21 |
| *Andrena bimaculata* | 0 | 0 | 0 | 0 | 0 | 2 | 0 | 7 |
| *Andrena congruens* | 0 | 0 | 0 | 0 | 2 | 0 | 0 | 3 |
| *Andrena ferrugineicrus* | 0 | 0 | 0 | 0 | 1 | 0 | 0 | 0 |
| *Andrena flavipes* | 0 | 0 | 0 | 0 | 8 | 38 | 0 | 8 |
| *Andrena fulva* | 0 | 0 | 0 | 0 | 6 | 24 | 0 | 11 |
| *Andrena granulosa* | 0 | 0 | 0 | 0 | 0 | 0 | 0 | 1 |
| *Andrena haemorrhoa* | 0 | 0 | 0 | 0 | 7 | 0 | 0 | 21 |
| *Andrena hispania* | 0 | 0 | 0 | 0 | 2 | 0 | 0 | 0 |
| *Andrena humilis* | 0 | 0 | 0 | 0 | 2 | 0 | 0 | 4 |
| *Andrena impressa* | 0 | 0 | 0 | 0 | 8 | 0 | 0 | 10 |
| *Andrena labialis* | 0 | 0 | 0 | 0 | 10 | 32 | 0 | 14 |
| *Andrena lepida* | 0 | 0 | 0 | 0 | 3 | 0 | 0 | 1 |
| *Andrena livens* | 0 | 0 | 0 | 0 | 4 | 0 | 0 | 11 |
| *Andrena nigroaenea* | 0 | 1 | 0 | 0 | 15 | 11 | 0 | 45 |
| *Andrena orbitalis* | 0 | 0 | 0 | 0 | 0 | 0 | 0 | 2 |
| *Andrena ovatula* | 0 | 0 | 0 | 0 | 1 | 0 | 0 | 0 |
| *Andrena paucisquama* | 0 | 0 | 0 | 0 | 0 | 0 | 0 | 2 |
| *Andrena pilipes* | 0 | 0 | 0 | 0 | 5 | 1 | 0 | 8 |

| | | | | | | | | |
|---|---|---|---|---|---|---|---|---|
| *Andrena rhenana* | 0 | 0 | 0 | 0 | 0 | 0 | 0 | 16 |
| *Andrena rhyssonota* | 0 | 0 | 0 | 0 | 1 | 0 | 0 | 0 |
| *Andrena sardoa* | 0 | 0 | 0 | 0 | 3 | 24 | 0 | 41 |
| *Andrena schencki* | 0 | 0 | 0 | 0 | 1 | 1 | 0 | 3 |
| *Andrena senecionis* | 0 | 0 | 0 | 0 | 0 | 0 | 0 | 4 |
| *Andrena synadelpha* | 0 | 0 | 0 | 0 | 0 | 0 | 0 | 9 |
| *Andrena thoracica* | 0 | 0 | 0 | 0 | 8 | 1 | 0 | 0 |
| *Andrena tibialis* | 0 | 0 | 0 | 0 | 1 | 3 | 0 | 22 |
| *Andrena trimmerana* | 0 | 0 | 0 | 0 | 17 | 35 | 0 | 20 |
| *Andrena vulpecula* | 0 | 0 | 0 | 0 | 2 | 1 | 0 | 0 |
| *Anthidiellum strigatum* | 0 | 0 | 0 | 0 | 0 | 0 | 0 | 10 |
| *Anthidium florentinum* | 0 | 0 | 0 | 0 | 0 | 0 | 19 | 29 |
| *Anthidium manicatum* | 0 | 0 | 0 | 0 | 0 | 0 | 1 | 15 |
| *Anthophora aestivalis* | 0 | 0 | 0 | 0 | 0 | 3 | 0 | 8 |
| *Anthophora affinis* | 0 | 0 | 0 | 0 | 0 | 1 | 0 | 0 |
| *Anthophora bimaculata* | 0 | 0 | 0 | 0 | 0 | 0 | 0 | 1 |
| *Anthophora crassipes* | 0 | 0 | 1 | 0 | 0 | 0 | 0 | 6 |
| *Anthophora dispar* | 0 | 0 | 3 | 2 | 0 | 11 | 0 | 12 |
| *Anthophora fulvodimidiata* | 0 | 0 | 0 | 0 | 0 | 0 | 0 | 5 |
| *Anthophora leucophaea* | 0 | 0 | 0 | 0 | 0 | 3 | 0 | 9 |
| *Anthophora plumipes* | 0 | 0 | 0 | 0 | 0 | 0 | 0 | 1 |
| *Anthophora retusa* | 0 | 0 | 0 | 0 | 0 | 2 | 0 | 6 |
| *Anthophora romandii* | 0 | 0 | 2 | 0 | 0 | 3 | 0 | 2 |
| *Ceratina chalybea* | 0 | 0 | 0 | 0 | 0 | 0 | 0 | 2 |
| *Ceratina cyanea* | 0 | 0 | 0 | 0 | 0 | 0 | 0 | 6 |
| *Ceratina gravidula* | 0 | 0 | 0 | 0 | 0 | 0 | 0 | 3 |
| *Ceratina mocsaryi* | 0 | 0 | 0 | 0 | 0 | 0 | 0 | 1 |
| *Coelioxys acanthura* | 0 | 0 | 0 | 0 | 0 | 0 | 0 | 1 |
| *Coelioxys brevis* | 0 | 0 | 0 | 0 | 0 | 0 | 0 | 1 |
| *Colletes cunicularius* | 0 | 0 | 0 | 0 | 5 | 29 | 0 | 24 |
| *Colletes hederae* | 0 | 0 | 0 | 0 | 0 | 0 | 0 | 14 |
| *Colletes nigricans* | 0 | 0 | 0 | 0 | 0 | 0 | 0 | 2 |
| *Colletes sierrensis* | 0 | 0 | 0 | 0 | 0 | 0 | 0 | 8 |
| *Dasypoda morotei* | 0 | 0 | 0 | 0 | 1 | 0 | 0 | 0 |
| *Epeolus fallax* | 0 | 0 | 0 | 0 | 0 | 0 | 0 | 3 |
| *Eucera caspica* | 0 | 0 | 0 | 1 | 0 | 0 | 0 | 0 |

| | | | | | | | | |
|---|---|---|---|---|---|---|---|---|
| *Eucera elongatula* | 0 | 0 | 0 | 0 | 0 | 4 | 0 | 28 |
| *Eucera longicornis* | 0 | 0 | 0 | 0 | 0 | 0 | 0 | 1 |
| *Eucera nigrilabris* | 0 | 0 | 0 | 0 | 0 | 0 | 0 | 1 |
| *Halictus fulvipes* | 0 | 0 | 0 | 0 | 0 | 0 | 0 | 1 |
| *Halictus gemmeus* | 0 | 0 | 0 | 0 | 0 | 0 | 0 | 2 |
| *Halictus quadricinctus* | 0 | 0 | 0 | 0 | 0 | 0 | 0 | 1 |
| *Halictus scabiosae* | 0 | 0 | 0 | 0 | 0 | 0 | 0 | 1 |
| *Halictus smaragdulus* | 0 | 0 | 0 | 0 | 0 | 0 | 0 | 5 |
| *Halictus subauratus* | 0 | 0 | 0 | 0 | 0 | 0 | 0 | 3 |
| *Heriades crenulatus* | 0 | 0 | 0 | 0 | 0 | 0 | 0 | 4 |
| *Hylaeus communis* | 0 | 0 | 0 | 0 | 0 | 0 | 0 | 4 |
| *Hylaeus gibbus* | 0 | 0 | 0 | 0 | 0 | 0 | 0 | 1 |
| *Hylaeus variegatus* | 0 | 0 | 0 | 0 | 0 | 0 | 0 | 7 |
| *Icteranthidium grohmanni* | 0 | 0 | 0 | 0 | 0 | 0 | 0 | 2 |
| *Icteranthidium laterale* | 0 | 0 | 0 | 0 | 0 | 0 | 0 | 1 |
| *Lasioglossum brevicorne* | 0 | 0 | 0 | 0 | 0 | 0 | 0 | 1 |
| *Lasioglossum buccale* | 0 | 0 | 0 | 0 | 0 | 0 | 0 | 1 |
| *Lasioglossum calceatum* | 0 | 0 | 0 | 0 | 0 | 0 | 0 | 1 |
| *Lasioglossum marginatum* | 0 | 0 | 0 | 0 | 0 | 0 | 0 | 5 |
| *Lasioglossum puncticolle* | 0 | 0 | 0 | 0 | 0 | 0 | 0 | 1 |
| *Lasioglossum villosulum* | 0 | 0 | 0 | 0 | 0 | 0 | 0 | 1 |
| *Lasioglossum xanthopus* | 0 | 0 | 0 | 0 | 0 | 0 | 0 | 2 |
| *Lithurgus chrysurus* | 0 | 0 | 0 | 0 | 0 | 0 | 0 | 9 |
| *Megachile albisecta* | 0 | 0 | 0 | 0 | 0 | 0 | 0 | 5 |
| *Megachile apicalis* | 0 | 0 | 0 | 0 | 0 | 0 | 0 | 1 |
| *Megachile centuncularis* | 0 | 0 | 0 | 0 | 0 | 0 | 0 | 1 |
| *Megachile giraudi* | 0 | 0 | 0 | 0 | 0 | 0 | 0 | 1 |
| *Megachile lagopoda* | 0 | 0 | 0 | 0 | 0 | 0 | 0 | 1 |
| *Megachile octosignata* | 0 | 0 | 0 | 0 | 0 | 0 | 0 | 1 |
| *Megachile pilicrus* | 0 | 0 | 0 | 0 | 0 | 0 | 0 | 2 |
| *Megachile pilidens* | 0 | 0 | 0 | 0 | 0 | 0 | 0 | 7 |
| *Megachile rotundata* | 0 | 0 | 0 | 0 | 0 | 0 | 0 | 1 |
| *Melecta albifrons* | 0 | 0 | 0 | 0 | 0 | 0 | 0 | 2 |
| *Nomada flavilabris* | 0 | 0 | 0 | 0 | 0 | 0 | 0 | 1 |
| *Nomada melathoracica* | 0 | 0 | 0 | 0 | 0 | 0 | 0 | 1 |
| *Nomada succincta* | 0 | 0 | 0 | 0 | 0 | 1 | 0 | 2 |

| | | | | | | | | |
|---|---|---|---|---|---|---|---|---|
| *Osmia bicornis* | 0 | 0 | 0 | 0 | 0 | 9 | 0 | 33 |
| *Osmia cornuta* | 0 | 0 | 0 | 0 | 0 | 15 | 0 | 13 |
| *Osmia niveata* | 0 | 0 | 0 | 0 | 0 | 0 | 0 | 1 |
| *Osmia tricornis* | 0 | 0 | 0 | 0 | 0 | 0 | 0 | 1 |
| *Panurgus banksianus* | 0 | 0 | 0 | 0 | 3 | 0 | 0 | 9 |
| *Panurgus perezi* | 0 | 0 | 0 | 0 | 0 | 0 | 0 | 2 |
| *Rhodanthidium sticticum* | 0 | 0 | 0 | 0 | 0 | 3 | 0 | 10 |
| *Thyreus ramosus* | 0 | 0 | 0 | 0 | 0 | 0 | 0 | 2 |
| *Thyreus truncatus* | 0 | 0 | 0 | 0 | 0 | 0 | 0 | 1 |
| *Xylocopa cantabrita* | 0 | 0 | 0 | 0 | 0 | 2 | 0 | 6 |
| *Xylocopa valga* | 0 | 0 | 1 | 0 | 0 | 5 | 0 | 5 |
| *Xylocopa violacea* | 1 | 0 | 1 | 0 | 0 | 3 | 0 | 27 |
