## Appendix S2 for "Body mass decline in a Mediterranean community of solitary bees supports the size shrinking effect of climatic warming"

**Appendix S2. Supporting information for analyses of temperature trends in the study region.**

FIGURE S1. Map showing the location of the 10 weather stations included in the analyses of daily maximum and minimum temperature trends (triangles) in relation to the locations where bees were sampled (dots).

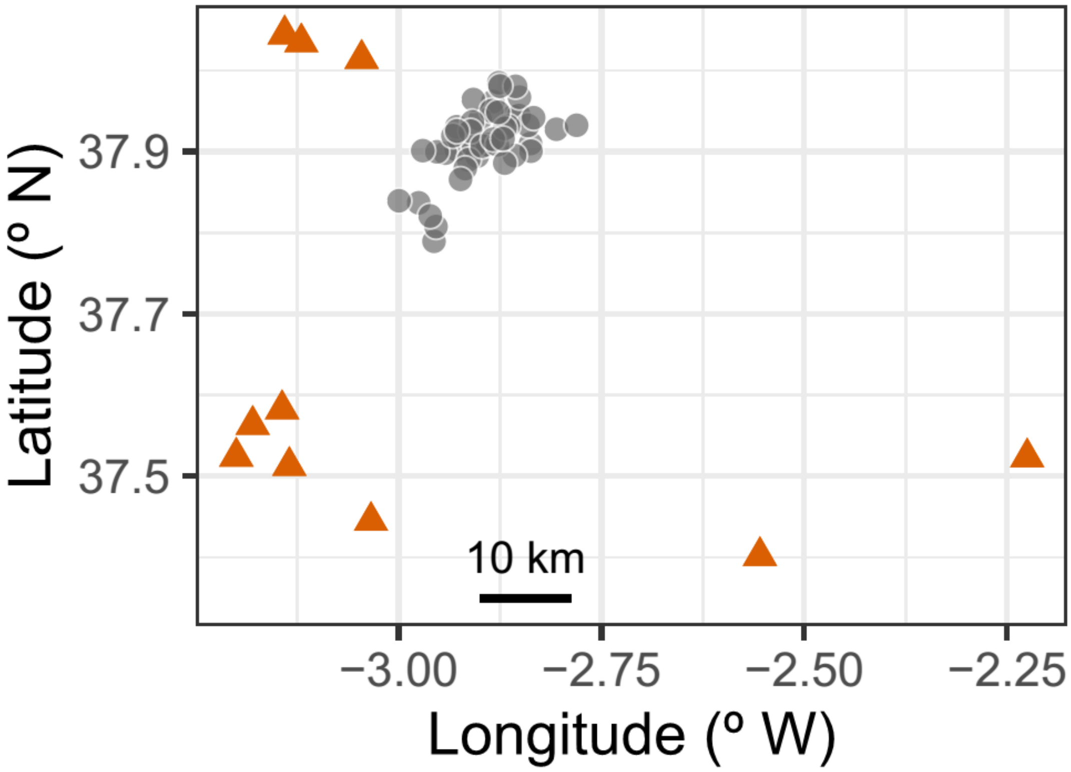

TABLE S1. Geographical information on the 10 weather stations whose data were included in the analyses of climatic change in the study region over 2000-2022, and summary of linear mixed models used to test for supra-annual linear trends in yearly means of daily maximum and minimum temperatures. Fixed-effect parameter estimates are the slopes of the fitted temperature/year linear relationships, and represent the rate of temperature change expressed in ºC·year$^{-1}$. Confidence intervals of parameter estimates were obtained by bootstraping each model with 1,000 iterations. To account for multiple testing, *P*-values shown were corrected using the Benjamini-Hochberg procedure.

| | | | | | Fixed-effect parameter | | Significance test | |
|---|---|---|---|---|---|---|---|---|
| Station | Elevation (m) | Latitude (º N) | Longitude (º W) | Temperature | Estimate | 95% Confidence interval | Chi-square | *P*-value |
| Huesa | 779 | 37.4450 | 3.0342 | Daily maximum | 0.0416 | 0.0286, 0.0557 | 36.39 | 3.2E-09 |
| | | | | Daily minimum | -0.0012 | -0.0106, 0.0080 | 0.06 | 0.84 |
| Jódar | 486 | 37.5242 | 3.2003 | Daily maximum | 0.0994 | 0.0775, 0.1205 | 85.62 | 7.3E-20 |
| | | | | Daily minimum | 0.0412 | 0.0227, 0.0591 | 21.24 | 6.2E-06 |
| Pozo Alcón | 881 | 37.4018 | 2.5548 | Daily maximum | 0.0485 | 0.0346, 0.0628 | 48.02 | 9.4E-12 |
| | | | | Daily minimum | -0.0057 | -0.0162, 0.0037 | 1.24 | 0.29 |

| | | | | | | | | |
|---|---|---|---|---|---|---|---|---|
| Puebla de Don Fadrique | 1017 | 37.5233 | 2.2254 | Daily maximum | 0.0496 | 0.0353, 0.0625 | 51.67 | 1.6E-12 |
| | | | | Daily minimum | 0.0185 | 0.0084, 0.0290 | 12.57 | 5.2E-04 |
| Sabiote | 791 | 38.0446 | 3.1407 | Daily maximum | 0.0645 | 0.0512, 0.0778 | 83.80 | 1.6E-19 |
| | | | | Daily minimum | 0.0193 | 0.0092, 0.0292 | 13.04 | 4.4E-04 |
| San José de los Propios | 494 | 37.5128 | 3.1349 | Daily maximum | 0.0692 | 0.0556, 0.0829 | 104.15 | 1.2E-23 |
| | | | | Daily minimum | -0.0007 | -0.0110, 0.0095 | 0.02 | 0.89 |
| Santo Tomé | 537 | 38.0145 | 3.0458 | Daily maximum | 0.0741 | 0.0587, 0.0881 | 99.43 | 1.0E-22 |
| | | | | Daily minimum | 0.0122 | 0.0028, 0.0215 | 6.00 | 0.018 |
| Torreperogil | 535 | 37.5827 | 3.1438 | Daily maximum | 0.0648 | 0.0451, 0.0839 | 35.91 | 3.8E-09 |
| | | | | Daily minimum | 0.0356 | 0.0207, 0.0505 | 22.91 | 2.8E-06 |
| Úbeda | 343 | 37.5634 | 3.1801 | Daily maximum | 0.0649 | 0.0510, 0.0775 | 91.75 | 3.9E-21 |
| | | | | Daily minimum | -0.0081 | -0.0202, 0.0029 | 1.85 | 0.20 |
| Villacarrillo | 649 | 38.0348 | 3.1201 | Daily maximum | 0.1145 | 0.0923, 0.1345 | 104.27 | 1.2E-23 |
| | | | | Daily minimum | 0.0876 | 0.0720, 0.1028 | 128.54 | 1.7E-28 |
