## Appendix S3 for "Body mass decline in a Mediterranean community of solitary bees supports the size shrinking effect of climatic warming"

### Appendix S3. Check of mixed model assumptions

FIGURE S1. Visual check of linearity, homocedasticity, noncollinearity, normality of residuals, and normality of random effects, for the mixed model testing the effect of sampling year on individual body mass of solitary bees.

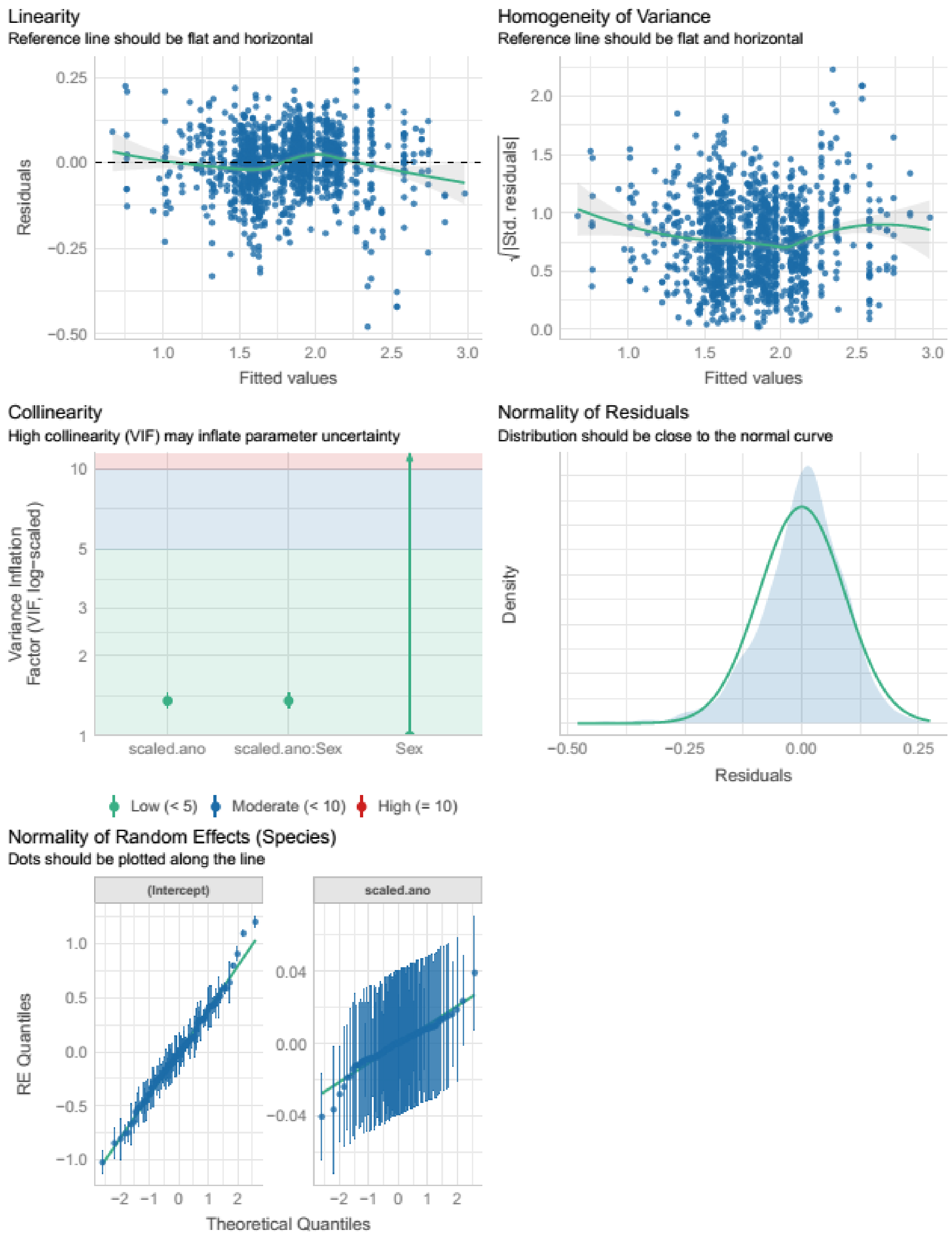
