## Appendix S4 for "Body mass decline in a Mediterranean community of solitary bees supports the size shrinking effect of climatic warming"

**Appendix S4. Paired comparisons of mean body mass for individual species.**

FIGURE S1. Paired within-species comparisons of mean body mass for the subset of 33 species in 10 genera which had males, females or both sexes sampled in the old (1990-1997) and recent (2022) periods. Each line connect the old and recent averages for the same species.

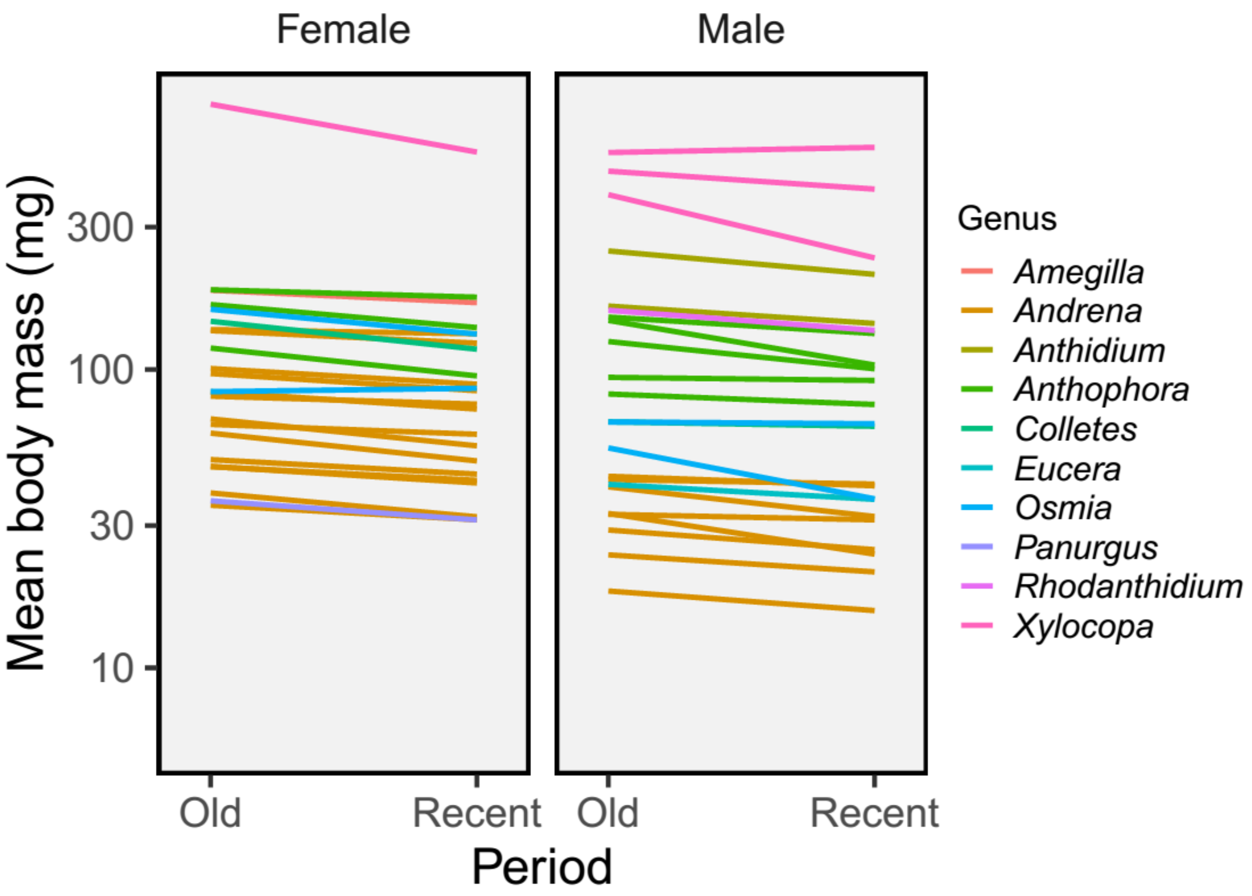
